# Accurate Diagnosis of Colorectal Cancer Based on Histopathology Images Using Artificial Intelligence

**DOI:** 10.1101/2020.03.15.992917

**Authors:** Kuan-Song Wang, Gang Yu, Chao Xu, Xiang-He Meng, Jianhua Zhou, Changli Zheng, Zhenghao Deng, Li Shang, Ruijie Liu, Shitong Su, Xunjian Zhou, Qingling Li, Juanni Li, Jing Wang, Kewen Ma, Jialin Qi, Zhenmin Hu, Ping Tang, Jeffrey Deng, Xiang Qiu, Bo-Yang Li, Wen-Di Shen, Ru-Ping Quan, Jun-Ting Yang, Lin-Ying Huang, Yao Xiao, Zhi-Chun Yang, Zhongming Li, Sheng-Chun Wang, Hongzheng Ren, Changjiang Liang, Wei Guo, Yanchun Li, Heng Xiao, Yonghong Gu, Jing-Ping Yun, Dan Huang, Zhigang Song, Xiangshan Fan, Ling Chen, Xiaochu Yan, Zhi Li, Zhong-Chao Huang, Jufang Huang, Joseph Luttrell, Chao-Yang Zhang, Weihua Zhou, Kun Zhang, Chunhui Yi, Hui Shen, Yu-Ping Wang, Hong-Mei Xiao, Hong-Wen Deng

## Abstract

**Background:** Accurate and robust pathological image analysis for colorectal cancer (CRC) diagnosis is time-consuming and knowledge-intensive, but is essential for CRC patients’ treatment. The current heavy workload of pathologists in clinics/hospitals may easily lead to unconscious misdiagnosis of CRC based on their daily image analyses.

**Methods:** Based on a state-of-the-art transfer-learned deep convolutional neural network in artificial intelligence (AI), we proposed a novel patch aggregation strategy for clinic CRC prediction/diagnosis using weakly labeled pathological whole slide image (WSI) patches. This approach was trained and validated using an unprecedented and enormously large number of 170,099 patches, >14,680 WSIs, from >9,631 subjects that covered diverse and representative clinical cases from multi-independent-sources across China, U.S., and Germany.

**Results:** Our innovative AI tool was consistently nearly perfectly agreed with (average Kappa-statistic 0.896) and even often better than most of the experienced expert pathologists when tested in diagnosing CRC WSIs from multi-centers. The average area under the receiver operating characteristics curve (AUC) of AI was greater than that of the pathologists (0.981 vs 0.970) and achieved the best performance among the application of other AI methods to CRC diagnosis. Our AI-generated heatmap highlights the image regions of cancer tissue/cells.

**Conclusions:** This first-ever generalizable AI system can handle large amounts of WSIs consistently and robustly without potential bias due to fatigue commonly experienced by clinical pathologists. Hence, it will drastically alleviate the heavy clinical burden of daily pathology diagnosis, and improve the treatment for CRC patients. This tool is generalizable to other cancer diagnosis based on image recognition.

## Introduction

Colorectal cancer (CRC) is the third leading cancer by incidence (6.1%) but second for mortality (9.2%) worldwide^1^. The global burden of CRC was expected to increase 60% by 2030, in terms of new cases and deaths^2^. The accurate and prompt detection of CRC is essential to improve treatment effectiveness and survivorship. The current diagnosis of CRC requires an extensive visual examination of digital whole slide images (WSIs) of the hematoxylin and eosin (H&E) stained specimens obtained from formalin-fixed paraffin-embedded (FFPE) or frozen tissues by highly specialized pathologists. The challenges for the WSI analysis include very large image size (>10,000×10,000 pixels), histological variations in size, shape, texture, and staining of nuclei, making the diagnosis complicated and time consuming^3^. In most modern pathology departments, the average consultative workload increases by ∼5-10% annually ^4^. The current trends clearly indicate a shortage of pathologists around the world, including U.S.A.^5^ and low-to middle-income countries^6^. The above situations will result in overwork for pathologists and may easily lead to higher chances of deficiencies in their routine work and dysfunctions of the pathology laboratories with more laboratory errors^4^. While the demands of colon specimen examination in gastroenterology clinics are high, the training time of pathologists is long (>10 years)^7^. It is thus imperative to develop reliable tools for pathological image analysis and CRC detection that can improve clinical efficiency and efficacy without unintended human bias during diagnosis.

State-of-the-art artificial intelligence (AI) approaches, such as deep learning (DL), are very powerful in classification and prediction. There have been many successful applications of DL, specifically convolutional neural network (CNN), in WSI analysis for cancers of, e.g., lung^8,9^, breast^10,11^, prostate^12^, and skin^13,14^. Most of the existing CNN for the CRC WSI analysis focused on the pathology work after cancer determination, including grade classification^15^, tumor cells detection and classification^16-18^, survivorship prediction^19-21^, etc. Although they resulted in reasonably high accuracy, their study sample sizes are limited and hence not fully represent the numerous histologic variants of CRC that have been defined, including tubular, mucinous, signet ring cell, and others^22^. The situation thus would inflate the prediction error when applied to different independent samples. Meanwhile, most of the current DL models were developed from single data source without thorough validation using independent data. They only calculated the accuracy of patches without diagnosing WSIs or the patients. Their general applicability for CRC WSI diagnosis in various clinical settings, which may involve heterogeneous platforms and image properties, remains unclear. A DL approach generalizable to daily pathological CRC diagnosis, to relieve clinical burden of pathologists, is yet to be developed.

Here, we developed a novel automated approach centered on weakly labeled supervised DL, a CNN using Inception-v3 architecture^23^ with weights initialized from transfer learning, for the very first clinical CRC diagnosis. Our work is based on WSIs from multiple independent hospitals/sources in China (8,554 patients), U.S. (1,077 patients), and Germany (>111 slides). Transfer learning is a highly effective and efficient DL technique for image analysis that can utilize previously learned knowledge on general images for medical images analyses^24^. Further, weakly labeled supervised learning, i.e. with only coarse-grained labels at image-level, is advantageous in training massive and diverse datasets without exact labelling at object-levels, such as the cancer cells^12^. This study has high practical value for improving the effectiveness and efficiency of CRC diagnosis and thus treatment. It highlights the general significance of the application of AI to image analyses of other types of cancers.

## Materials and Methods

### Colorectal cancer whole-slide image dataset

We collected 14,234 CRC WSIs from fourteen independent sources (Table 1). All data were de-identified. The largest image set was from 6,876 patients admitted between 2010 and 2018 in Xiangya Hospital (XH), Central South University (CSU, Changsha, China). XH is the largest hospital in Hunan Province and was established in 1906 with a close affiliation with Yale University^25^. The other independent sources were TCGA of US (https://portal.gdc.cancer.gov/)^26^, NCT-UMM of Germany (https://zenodo.org/record/1214456#.XgaR00dTm00)^20^, Adicon Clinical Laboratories, INC (ACL), and eleven hospitals in China (detailed in Table 1). The hospitals involved are located in the major metropolitan areas of China serving >139 million population, including those most prestigious hospitals in pathology in China: XH, FUS, CGH, SWH, and AMU; other state-level esteemed hospitals: SYU, NJD, GPH, HPH, and TXH; and a regional reputable PCH. All WSIs were from FFPE tissues, except part of TCGA WSIs were from frozen tissues^27^. The process of collection, quality control, and digitalization of the WSIs was described in Supplementary-Text 1.a.

**Table 1.** Usage of datasets from multi-center data source.

| Data source | Dataset Usage | Sample preparation | Examination type<br>Radical surgery /<br>Colonoscopy | Population * | CRC |  | Non-CRC |  | Total |  |
| --- | --- | --- | --- | --- | --- | --- | --- | --- | --- | --- |
|  |  |  |  |  | subjects | slides | subjects | slides | subjects | slides |
| Xiangya Hospital (XH) | A | FFPE | 100% / 0% | Changsha, China | 614 | 614 | 228 | 228 | 842 | 842 |
| NCT-UMM (NCT-CRC-HE-100K) | B | FFPE | NA | Germany | NA | NA | NA | NA | NA | 86 |
| NCT-UMM (CRC-VAL-HE-7K) | B | FFPE | NA | Germany | NA | NA | NA | NA | NA | 25 |
| XH | C | FFPE | 80% / 20% | Changsha, China | 3,990 | 7,871 | 1,849 | 2,132 | 5,839 | 10,003 |
| XH | D | FFPE | 89% / 11% | Changsha, China | 98 | 99 | 97 | 114 | 195 | 213 |
| Pingkuang Collaborative Hospital (PCH) | C & D | FFPE | 60% / 40% | Jiangxi, China | 50 | 50 | 46 | 46 | 96 | 96 |
| The Third Xiangya Hospital of CSU (TXH) | C & D | FFPE | 61% / 39% | Changsha, China | 48 | 70 | 48 | 65 | 96 | 135 |
| Hunan Provincial People's Hospital (HPH) | C & D | FFPE | 61% / 39% | Changsha, China | 49 | 50 | 49 | 49 | 98 | 99 |
| ACL | C & D | FFPE | 22% / 78% | Changsha, China | 100 | 100 | 107 | 107 | 207 | 207 |
| Fudan University Shanghai Cancer Center (FUS) | C & D | FFPE | 97% / 3% | Shanghai, China | 100 | 100 | 98 | 98 | 198 | 198 |
| Guangdong Provincial People's Hospital (GPH) | C & D | FFPE | 77% / 23% | Guangzhou, China | 100 | 100 | 85 | 85 | 185 | 185 |
| Nanjing Drum Tower Hospital (NJD) | C & D | FFPE | 96% / 4% | Nanjing, China | 100 | 100 | 97 | 97 | 197 | 197 |
| Southwest Hospital (SWH) | C & D | FFPE | 93% / 7% | Chongqing, China | 99 | 99 | 100 | 100 | 199 | 199 |
| The First Affiliated Hospital Air Force Medical University (AMU) | C & D | FFPE | 95% / 5% | Xi'an, China | 101 | 101 | 104 | 104 | 205 | 205 |
| Sun Yat-Sen University Cancer Center (SYU) | C & D | FFPE | 100% / 0% | Guangzhou, China | 91 | 91 | 6 | 6 | 97 | 97 |
| Chinese PLA General Hospital (CGH) | C | FFPE | NA | Beijing, China | 0 | 0 | 100 | 100 | 100 | 100 |
| TCGA (TCGA-Frozen) | C | Frozen | 100% / 0% | U.S. | 631 | 1214 | 110 | 133 | 631** | 1347 |
| TCGA (TCGA-FFPE) | C | FFPE | 100% / 0% | U.S. | 441 | 441 | 5 | 5 | 446 | 446 |
| **Total** |  |  |  |  | 6,612 | 11,100 | 3,129 | 3,469 | 9,631 | 14,680 |

\*Location map available in Supplementary Text 1.a. \*\* For the TCGA –Frozen data only, the non-CRC slides were made with normal intestinal tissues on part of the CRC slides.

We formed four datasets (Table 1). Dataset-A with slides from XH only was used for patch-level training and testing (Table 2). We carefully selected WSIs to include all common tumor histological subtypes. Pathologists weakly labeled the patches from WSIs as either containing or not cancer cells/tissues without complete information of cancer cells/tissues (location, shape, demarcation etc.). Dataset-B consisting of two independent subsets in NCT-UMM were used for patch-level external validation. The overall split for patch-level training, testing, and external validation was about 2:1:5. Dataset-C used for patient-level validation was composed of slides from XH, the other hospitals, ACL, and frozen and FFPE samples of TCGA. Given the highly imbalance of cancer and non-cancer slides in SYU and CGH (Table 1), they were combined in Dataset-C. Dataset-D used for the Human-AI contest contained approximately equal number of slides from XH, the other hospitals, and ACL. Supplementary-Text 1.b summarized the allocation of slides in the different datasets, sampling strategy, and other details.

**Table 2.** Dataset-A (training and testing) and Dataset-B (external validation) for patch-level analysis.

| **Dataset** | **Cancer** |  |  | **Non-cancer** |  |  | **Total** |  |  |
| --- | --- | --- | --- | --- | --- | --- | --- | --- | --- |
|  | subjects | slides | patches | subjects | slides | patches | subjects | slides | patches |
| **Training** | 406 | 406 | 19,940 | 153 | 153 | 22,715 | 559 | 559 | 42,655 |
| **Testing** | 208 | 208 | 10,116 | 75 | 75 | 10,148 | 283 | 283 | 20,264 |
| **Validation** | NA | NA | 15,550 | NA | NA | 91,630 | NA | 111 | 107,180* |
| **Total** | >614 | >614 | 45,606 | >228 | >228 | 124,493 | >842 | 953 | 170,099 |

\* There are two datasets used for validation. The number is the sum of the two datasets.

Our approach to predict patient cancerous status involved two steps: DL prediction for local patches and patch-level results aggregation for patient-level diagnosis (Supplementary-Figure 1).

**Figure 1.**
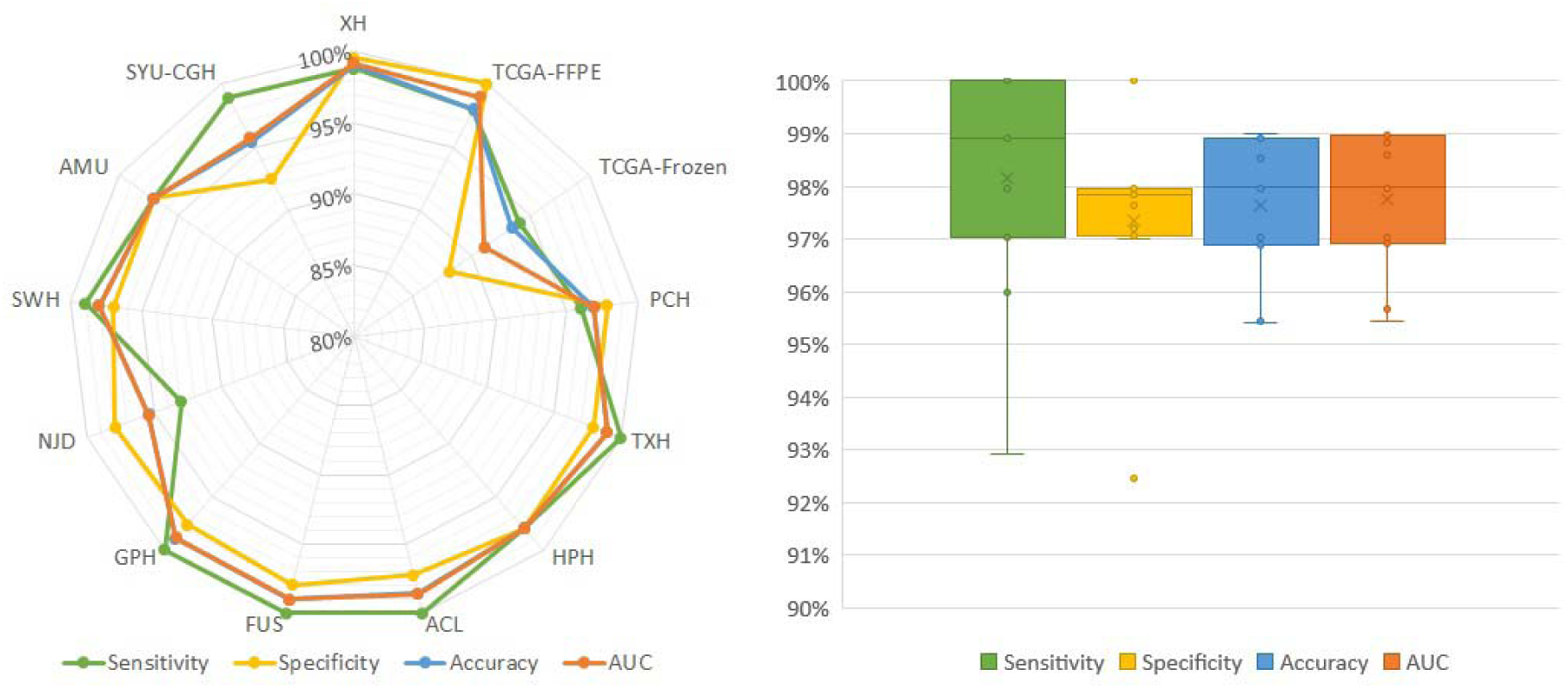
Patient-level testing performance on twelve independent datasets from Dataset-C. Left: the radar map of the sensitivity, specificity, accuracy, and AUC in each dataset from Dataset-C. Right: the boxplot showing the distribution of sensitivity, specificity, accuracy, and AUC in datasets (excluding XH and TCGA-Frozen). The horizontal bar in the box indicates the median, while the cross indicates the mean. Circles represent data points.

### Patch-level training by deep learning

Our DL model used Inception-v3 as the CNN architecture to classify cancerous and normal patches. The Inception network using different kernel sizes is specifically powerful in learning diagnostic information in pathological image from differing scales and has achieved near human expert performance in the analyses of other cancer types^8,13,28,29^. There are a few Inception architectures performed well on the ImageNet dataset^30^ and WSIs analysis^30^, such as the Inception-v1^31^, Inception-v3^23^, Inception-v4^32^. We chose Inception-v3 based on extensive comparison of their patch-level and patient-level performance in testing sets. (Supplementary-Text 1.c).

We initialized the CNN by transfer learning with pre-trained weights from ImageNet^23^, which were optimized to capture the structures in general images^24^. With transfer learning, our model can recognize pivotal image features for CRC diagnosis most efficiently. After preprocessing, the cropped non-overlapping patches from each WSI in training set were fed into the initialized CNN for fine-tuning (Supplementary-Text 1.d).

### Patient diagnosis and false positive control

Considering the high FPR (false positive rate) accumulated from multiple patch-level predictions, we proposed a novel patch-cluster-based aggregation method for slide-level prediction based on the fact that the tumor cells tend to gather together (especially at 20× magnification). Motivated by the clustering inference of fMRI^33^, we predicted the WSI as cancer positive if there were several positive patches topologically connected as a cluster on the slide (defined by the cluster size), such as four patches as a square. Otherwise, we predicted the slide as negative. We tested various cluster sizes and picked four due to an empirically observed best balance of sensitivity and FPR in the testing dataset (Supplementary-Text 1.e). We provided the patient-level diagnosis combining results from (the union of) all of the patient’s slides. So that the patient will be diagnosed as having cancer as long as one of the slides is diagnosed so.

### Human-AI contest

Six pathologists (A-F) with varying experience of 1 to 18 clinical practice years joined the contest (Supplementary-Table 1). The pathologists independently provided a diagnosis specifying cancer or non-cancer for each patient after reading the WSIs in Dataset-D. None of them participated in the data collection or labeling. An independent analyst blindly summarized and compared the accuracy and speed of AI and human experts in performing diagnosis. Details of the statistical methods are in Supplementary-Text 1.f.

## Results

### Highest accuracies in patch-level prediction by our model

We divided the 842 WSIs from Dataset-A (Table 2) into 62,919 non-overlapping patches to construct the CNN for patch-level prediction based on fine-tuning of Inception-v3. An average of ∼75 patches per WSI were included to ensure an appropriate and comprehensive representation of cancer and normal tissue characteristics. Three major CRC histological subtypes were involved for the training and testing, including 74.76% tubular, 24.59% mucinous, and 0.65% signet ring cell patches, roughly reflecting their clinical incidences^34^. In the training, 19,940 (46.75%) patches had cancer, and 22,715 (53.25%) patches were normal. Using another independent set of 10,116 (49.92%) cancer and 10,148 (50.08%) non-cancer patches, the AI for patch-level prediction achieved a testing accuracy of 98.11% and an AUC of 99.83%. The AUC outperformed that of all the previous AI studies for CRC diagnosis and prediction (79.2%-99.4%) and even for the majority of other types of cancer (82.9%-99.9%, Supplementary-Table 2). The specificity was 99.22% and the sensitivity 96.99%, both outstanding. In the external validation Dataset-B, our model yielded an accuracy and AUC of 96.07% and 98.32% in NCT-CRC-HE-100K, and 94.76% and 98.45% in CRC-VAL-HE-7K, which matched the performance from in-house data and outplayed the patch-level validation analysis in other AI studies (AUC 69.3%-95.0%, Supplementary-Table 2).

### Diagnosis of CRC at patient level using DL-predicted patches

Our AI approach was tested for patient diagnosis with 13,514 slides from 8,594 patients (Dataset-C). In the largest subset (5,839 patients) from XH, our approach produced an accuracy of 99.02% and an AUC of 99.16% (Figure 1, Supplementary-Table 3). In other independent multi-center datasets, our approach consistently performed very well. For the FFPE slides from other hospitals, TCGA-FFPE, and ACL, the AI approach yielded an average AUC and accuracy higher than 97.65% (Figure 1). For frozen slides TCGA-Frozen, the AI accuracy and AUC were 93.44% and 91.05% respectively (Figure 1). Our AUC values (ranging from 91.05% to 99.16%) were higher than that of other AI-based approaches for independent datasets (ranging from 83.3% to 94.1%), while the majority of those earlier AI approaches were tested on datasets of much smaller sample sizes (Supplementary-Table 2). The limited number of negative slides in TCGA may result in an imbalanced classification problem that needs further investigation, which is beyond the scope of this study. The results on TCGA-Frozen slides showed that our method did learn the histological morphology of cancer and normal tissues for cancer diagnosis, which is preserved in both the FFPE and frozen samples, even though our method was developed based on the FFPE samples. See Supplementary-Table 3 for complete patient-level result.

### Contest with six human experts

The performance of our AI approach was consistently comparable to the pathologists in diagnosing 1,831 WSIs from independent centers (Dataset-D, Figure 2). The AI resulted in an overall accuracy and AUC of 98.06% and 98.10%, which both ranked third out of the seven competitors (AI plus the six pathologists) and were greater than the average of the pathologists (accuracy 97.14% and AUC 96.95%). The AI yielded the highest sensitivity (98.16%) relative to the average (97.47%) of the pathologists (Supplementary-Table 4). The pathologists (D and E) who slightly outperformed the AI have 7 and 12 years of clinical experience respectively, while the AI outperformed the other 4 pathologists with 1, 3, 5, and 18 years of experience respectively. Cohen’s Kappa statistic (*K*) showed an excellent agreement (*K*≥0.858) between AI and every pathologist (Supplementary-Table 5). Our approach is thus proven generalizable to provide diagnosis support for potential CRC subjects like an independent pathologist, which can drastically relieve the heavy clinical burden and training cost of professional pathologists. Details of the Human-AI contest are given in Supplementary-Tables 3&4.

**Figure 2.**
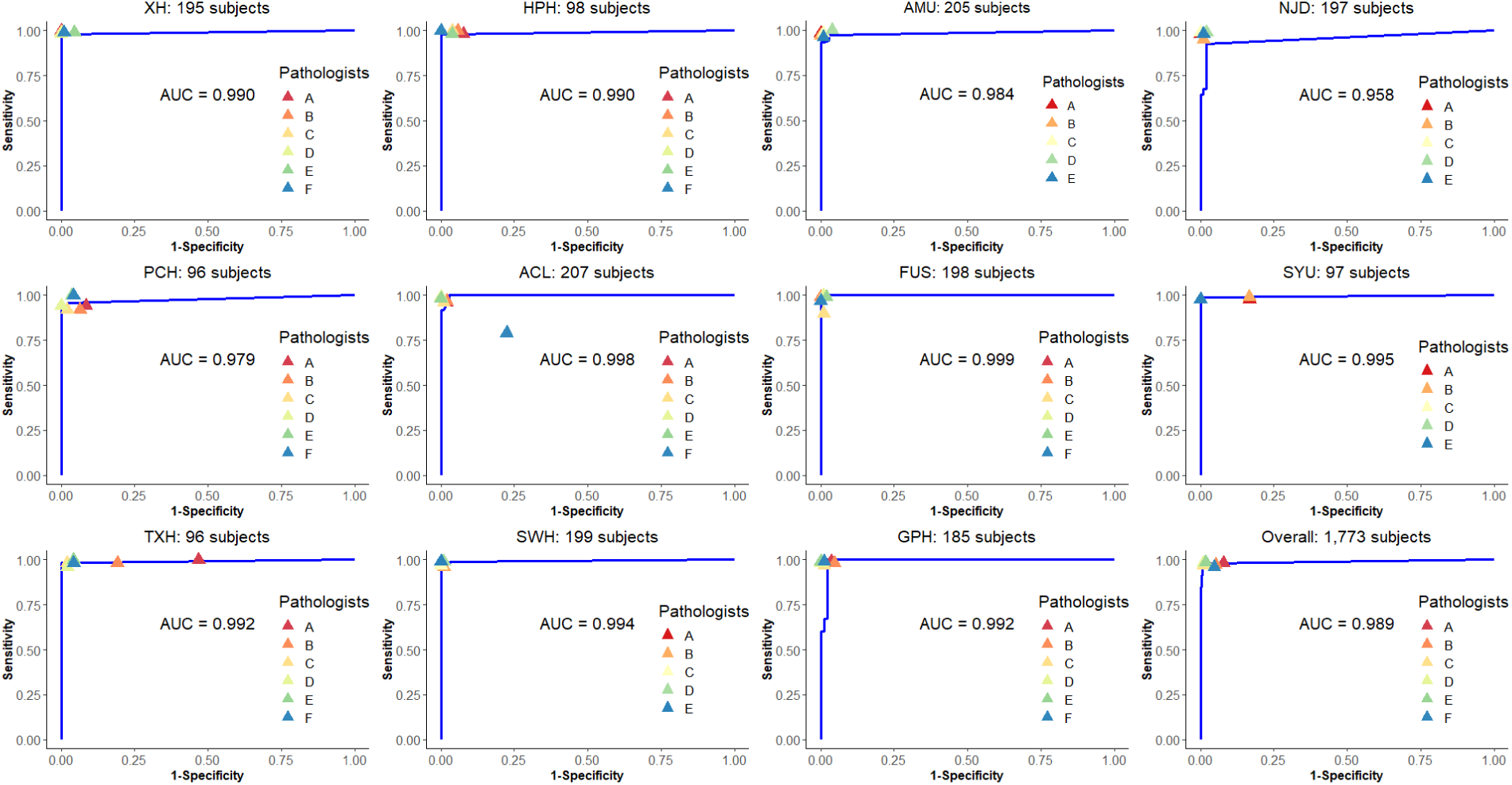
ROC analysis of AI and pathologists in the Human-AI contest using Dataset-D. The blue line was the estimated ROC curve for AI. The colored triangles indicate the sensitivity and specificity achieved by the six pathologists.

The pathologists were all informed to compete with our AI and with each other; hence, their performances were achieved under their best possible conditions with very best effort, which represented their highest skill with least error and fastest speed. However, with heavy workload in clinic, their performance in terms of accuracy and speed will not be as stable as that of AI. The current study of AI in cancer diagnosis using WSI have shown that AI can accurately diagnose in ∼20 seconds^8^ or less (∼13 seconds in our case). With evolved DL techniques and advanced computing hardware, the AI can constantly improve and provide steady, swift, and accurate first diagnosis for CRC or other cancers.

### Slide-level heatmap

Our approach offers an additional distinct feature: heatmap for highlighting potential cancer regions (as patches) in WSI. In Figure 3, we presented two WSIs, which were overlaid with the predicted heatmap. For both radical surgery WSI and colonoscopy WSI, the true cancerous region was highly overlapped with highlighted patches obtained by AI, which was also verified by pathologists. See more examples in Supplementary-Figure 2. In addition, to visualize informative regions utilized by DL for the CRC detection, we provided the activation maps in Supplementary-Figure 3.

**Figure 3.**
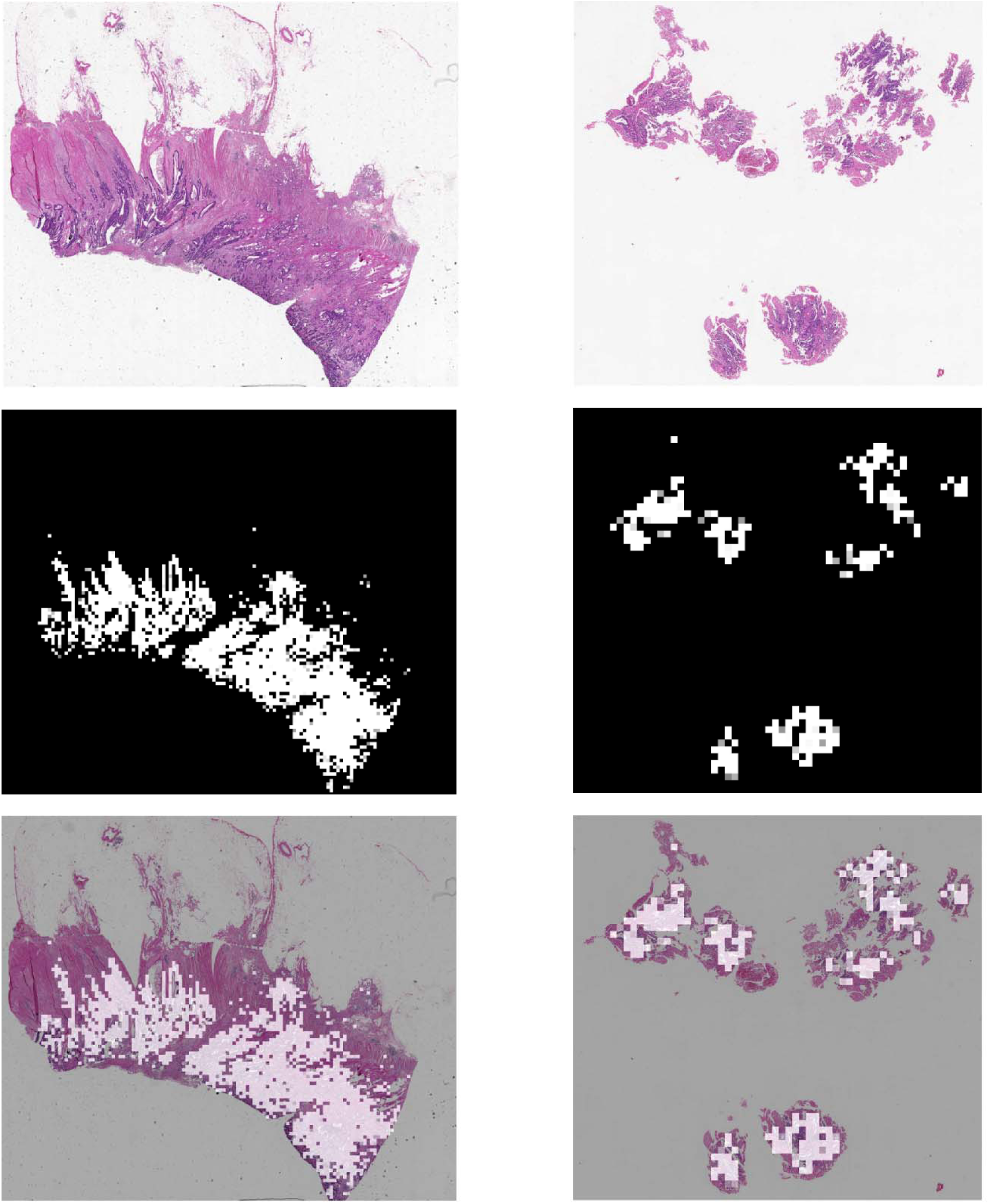
Heatmap produced by AI. Top row: WSI from radical surgery (left) and colonoscopy (right); middle row: AI predicted heatmap corresponding to the first row, with white coloration indicating predicted cancer region; bottom row: heatmap overlaid on the corresponding WSI.

## Discussion

We collected high-quality, comprehensive, and multiple independent human WSI datasets for training, testing, and external validation of our AI-based approach focusing on pathological diagnosis of CRC under common clinical settings. We mimicked the clinical procedure of WSI analysis, including the image digitalization, slide review, and expert consultations of the disputed slides. Different from other studies^18^, we did not apply any manual selection of slides or the area of interest when building the study dataset. Given the complex histologic variants of CRC, we randomly selected training patches from three most commonly seen subtypes roughly proportional to their incidences. The number of patches from images with large and small cancer tissue area was balanced and well represented in patch-level analysis. The collected images were labeled by agreement of at least two senior experts in CRC pathology (Supplementary text 1.b). The testing dataset from different locations in China, U.S., and Germany served as a representative pool for validation and generalization. Our dataset well represents the slides seen in clinics. Consequently, the trained AI model is robust and generalizable to analyze images of different production protocols and image quality.

For a fast-growing area, we are aware of that several new CNN architectures have been proposed after the completion of the study of the present paper, such as the DenseNet^35^, Squeeze-and-Excitation network^36^, ResNeXt^37^, etc. Although they have been shown to increase the prediction accuracy on ImageNet dataset compared to the Inception-v3, their performance on pathology images analysis and cancer diagnosis may await more focused technical comparison research.

There are several histological types that were too rare (less than 0.5% in incidence^38^) to be included, such as medullary, micropapillary, serrated. Our AI approach performed only slightly less satisfactory in frozen samples than in FFPE samples. With WSIs from rare types and more frozen samples available for training in the future, we expect our approach can be constantly improved to be more generalizable.

Most of the previous studies obtained the patient’s diagnosis by integrating the patch-level recognition results, since it is not feasible to process the large-size WSI directly. This strategy is difficult to control the accumulated FPR from multiple predictions based on individual patches. Recently, Coudray et al. used the proportion of positive patches or the average probability of all patches as the prediction criterion for the WSI^8^. Although their results were verified in three independent datasets (all with small sample sizes (340 slides)), their aggregation method may not be valid for those images with only a small area of cancer tissues where it will yield false negative findings for cancer patients. Instead, we proposed a novel aggregation strategy for patch-based WSI or patient-level prediction, which is intuitive and can easily balance the sensitivity and specificity. In practice, setting the cluster size to four is most likely to exceed the average accuracy of pathologists, while cluster size of two can be used for pathological screening with an average sensitivity of ∼99.78% and an average specificity of ∼72.29% according to our test data (Supplementary-Text 1.e).

Here, we developed a novel DL-based histopathological image classification approach for CRC diagnosis with the best performance achieved with the largest number of sample sizes and data sources in the field so far. Our approach was able to quickly and accurately distinguish CRC cases from healthy or inflammatory cases and was comparable to or even superior to pathologists in the testing of large-scale multi-center data. To the best of our knowledge, this is the first AI study for a reliable, generalized, and robust auxiliary tool for daily clinical pathology diagnosis of CRC initial screening. Our approach may also be adapted and applied to the histological analysis of other cancer types via the code available upon request.

## Acknowledgment

H.S. and H.W.D. were partially supported by grants from National Institutes of Health (R01AR059781, P20GM109036, R01MH107354, R01MH104680, R01GM109068, R01AR069055, U19AG055373, R01DK115679), the Edward G. Schlieder Endowment and the Drs. W. C. Tsai and P. T. Kung Professorship in Biostatistics from Tulane University. H.M.X was partially supported by the National Key Research and Development Plan of China (2017YFC1001103, 2016YFC1201805), National Natural Science Foundation of China (#81471453), and Jiangwang Educational Endowment. K.S.W was partially supported by the National Natural Science Foundation of China (#81673491) and the Natural Science Foundation of Hunan Province (#2015JJ2150). K.Z. was partially supported by grants from National Institutes of Health (2U54MD007595). Part of the computing for this project was performed at the OU Supercomputing Center for Education & Research (OSCER) at the University of Oklahoma (OU).

## Reference

1. Bray F, Ferlay J, Soerjomataram I, Siegel RL, Torre LA, Jemal A. Global cancer statistics 2018: GLOBOCAN estimates of incidence and mortality worldwide for 36 cancers in 185 countries. CA Cancer J Clin 2018.

2. Arnold M, Sierra MS, Laversanne M, Soerjomataram I, Jemal A, Bray F. Global patterns and trends in colorectal cancer incidence and mortality. Gut 2017;66:683–91.

3. Komura D, Ishikawa S. Machine Learning Methods for Histopathological Image Analysis. Comput Struct Biotechnol J 2018;16:34–42.

4. Maung R. Pathologists’ workload and patient safety. Diagnostic Histopathology 2016;22:283–7.

5. Metter DM, Colgan TJ, Leung ST, Timmons CF, Park JY. Trends in the US and Canadian Pathologist Workforces From 2007 to 2017. JAMA Netw Open 2019;2:e194337.

6. Sayed S, Lukande R, Fleming KA. Providing Pathology Support in Low-Income Countries. J Glob Oncol 2015;1:3–6.

7. Black-Schaffer WS, Morrow JS, Prystowsky MB, Steinberg JJ. Training Pathology Residents to Practice 21st Century Medicine: A Proposal. Acad Pathol 2016;3:2374289516665393.

8. Coudray N, Ocampo PS, Sakellaropoulos T, et al. Classification and mutation prediction from non-small cell lung cancer histopathology images using deep learning. Nat Med 2018.

9. Hua K-L, Hsu C-H, Hidayati SC, Cheng W-H, Chen Y-J. Computer-aided classification of lung nodules on computed tomography images via deep learning technique. OncoTargets and therapy 2015;8:2015–22.

10. Veta M, van Diest PJ, Willems SM, et al. Assessment of algorithms for mitosis detection in breast cancer histopathology images. Med Image Anal 2015;20:237–48.

11. Ehteshami Bejnordi B, Veta M, Johannes van Diest P, et al. Diagnostic Assessment of Deep Learning Algorithms for Detection of Lymph Node Metastases in Women With Breast Cancer. JAMA 2017;318:2199–210.

12. Campanella G, Hanna MG, Geneslaw L, et al. Clinical-grade computational pathology using weakly supervised deep learning on whole slide images. Nature Medicine 2019.

13. Esteva A, Kuprel B, Novoa RA, et al. Dermatologist-level classification of skin cancer with deep neural networks. Nature 2017;542:115–8.

14. Yu L, Chen H, Dou Q, Qin J, Heng PA. Automated Melanoma Recognition in Dermoscopy Images via Very Deep Residual Networks. IEEE Trans Med Imaging 2017;36:994–1004.

15. Sari CT, Gunduz-Demir C. Unsupervised Feature Extraction via Deep Learning for Histopathological Classification of Colon Tissue Images. IEEE Trans Med Imaging 2019;38:1139–49.

16. Sirinukunwattana K, Ahmed Raza SE, Yee-Wah T, Snead DR, Cree IA, Rajpoot NM. Locality Sensitive Deep Learning for Detection and Classification of Nuclei in Routine Colon Cancer Histology Images. IEEE Trans Med Imaging 2016;35:1196–206.

17. Haj-Hassan H, Chaddad A, Harkouss Y, Desrosiers C, Toews M, Tanougast C. Classifications of Multispectral Colorectal Cancer Tissues Using Convolution Neural Network. J Pathol Inform 2017;8:1.

18. Chaddad A, Tanougast C. Texture Analysis of Abnormal Cell Images for Predicting the Continuum of Colorectal Cancer. Anal Cell Pathol (Amst) 2017;2017:8428102.

19. Bychkov D, Linder N, Turkki R, et al. Deep learning based tissue analysis predicts outcome in colorectal cancer. Sci Rep 2018;8:3395.

20. Kather JN, Krisam J, Charoentong P, et al. Predicting survival from colorectal cancer histology slides using deep learning: A retrospective multicenter study. PLOS Medicine 2019;16:e1002730.

21. Skrede OJ, De Raedt S, Kleppe A, et al. Deep learning for prediction of colorectal cancer outcome: a discovery and validation study. Lancet 2020;395:350–60.

22. Fleming M, Ravula S, Tatishchev SF, Wang HL. Colorectal carcinoma: Pathologic aspects. J Gastrointest Oncol 2012;3:153–73.

23. Szegedy C, Vanhoucke V, Ioffe S, Shlens J, Wojna Z. Rethinking the Inception Architecture for Computer Vision. 2016 IEEE Conference on Computer Vision and Pattern Recognition (CVPR); 2016 27-30 June 2016. p. 2818–26.

24. Kermany DS, Goldbaum M, Cai W, et al. Identifying Medical Diagnoses and Treatable Diseases by Image-Based Deep Learning. Cell 2018;172:1122–31 e9.

25. Li LJ, Lu GX. How medical ethical principles are applied in treatment with artificial insemination by donors (AID) in Hunan, China: effective practice at the Reproductive and Genetic Hospital of CITIC-Xiangya. J Med Ethics 2005;31:333–7.

26. Grossman RL, Heath AP, Ferretti V, et al. Toward a Shared Vision for Cancer Genomic Data. N Engl J Med 2016;375:1109–12.

27. Cooper LA, Demicco EG, Saltz JH, Powell RT, Rao A, Lazar AJ. PanCancer insights from The Cancer Genome Atlas: the pathologist’s perspective. The Journal of pathology 2018;244:512–24.

28. Gulshan V, Peng L, Coram M, et al. Development and Validation of a Deep Learning Algorithm for Detection of Diabetic Retinopathy in Retinal Fundus PhotographsAccuracy of a Deep Learning Algorithm for Detection of Diabetic RetinopathyAccuracy of a Deep Learning Algorithm for Detection of Diabetic Retinopathy. JAMA 2016;316:2402–10.

29. Litjens G, Kooi T, Bejnordi BE, et al. A survey on deep learning in medical image analysis. Med Image Anal 2017;42:60–88.

30. Khosravi P, Kazemi E, Imielinski M, Elemento O, Hajirasouliha I. Deep Convolutional Neural Networks Enable Discrimination of Heterogeneous Digital Pathology Images. EBioMedicine 2018;27:317–28.

31. Szegedy C, Wei L, Yangqing J, et al. Going deeper with convolutions. 2015 IEEE Conference on Computer Vision and Pattern Recognition (CVPR); 2015 7-12 June 2015. p. 1–9.

32. Szegedy C, Ioffe S, Vanhoucke V, Alemi AA. Inception-v4, inception-resnet and the impact of residual connections on learning. Thirty-First AAAI Conference on Artificial Intelligence; 2017.

33. Heller R, Stanley D, Yekutieli D, Rubin N, Benjamini Y. Cluster-based analysis of FMRI data. Neuroimage 2006;33:599–608.

34. Liu T. Diagnostic Pathology. 3 ed. Beijing: People’s Medical Publishing House; 2013.

35. Huang G, Liu Z, Maaten Lvd, Weinberger KQ. Densely Connected Convolutional Networks. 2017 IEEE Conference on Computer Vision and Pattern Recognition (CVPR); 2017 21-26 July 2017. p. 2261–9.

36. Hu J, Shen L, Sun G. Squeeze-and-Excitation Networks. 2018 IEEE/CVF Conference on Computer Vision and Pattern Recognition; 2018 18-23 June 2018. p. 7132–41.

37. Veit A, Alldrin N, Chechik G, Krasin I, Gupta A, Belongie S. Learning from Noisy Large-Scale Datasets with Minimal Supervision. 2017 IEEE Conference on Computer Vision and Pattern Recognition (CVPR); 2017 21-26 July 2017. p. 6575–83.

38. Bosman FT, Carneiro F, Hruban RH, Theise ND. WHO classification of tumours of the digestive system. 4 ed. Lyon, France: International Agency for Research on Cancer; 2010.

