## Supplementary-Text for "Accurate Diagnosis of Colorectal Cancer Based on Histopathology Images Using Artificial Intelligence"

**Supplementary Files**

**1. Supplementary Text**

**1.a Collection and digitalization of the WSIs	2**

**1.b Dataset-A, B, C and D	3**

**1.c Comparison of different architectures at patch-level	7**

**1.d Image preprocessing and CNN training at patch-level	9**

**1.e Comparison of different cluster sizes for aggregation of patch-level results	11**

**1.f Statistical analysis and visualization	13**

**2. Supplementary Tables	14**

**3. Supplementary Figures	19**

**4. Reference	22**

**1.a Collection and digitalization of the WSIs**

After the data quality check introduced in the following paragraph, we collected more than 14,234 CRC WSIs from >9,185 patients from fourteen independent sources in China, U.S., and Germany (Figure S1), including: eleven hospitals (XH, TXH, HPH, PCH, FUS, AMU, NJD, GPH, CGH, SWH, SYU), a professional clinical service laboratory (ACL), and two public database (TCGA and NCT-UMM). The number of selected patients collected on the same day was limited to less than 50 to avoid potential over-representation of WSIs prepared on that day for subsequent analyses. The personnel who completed the random selection of WSIs did not participate in subsequent research.

According to the selected pathology ID, the technicians of the pathology department obtained the slides from the pathology archive library. After simply wiping off the dust on the surface, the slides were scanned by a KF-PRO-005 scanner (KFBIO company, Ningbo, China) at a 20× magnification. The scanning speed was ~40 seconds per slide. A quick visual examination of image quality was conducted to ensure the shape and location of tissue/cells on the digital slide was clear. Other factors, such as color differences, dirt in the background, and sharpness at the edge of cells were not considered. At this step, we removed 249 slides from 141 XH subjects and 26 slide images from TCGA that were low-resolution, unclear, obscure, or contained no tissues.

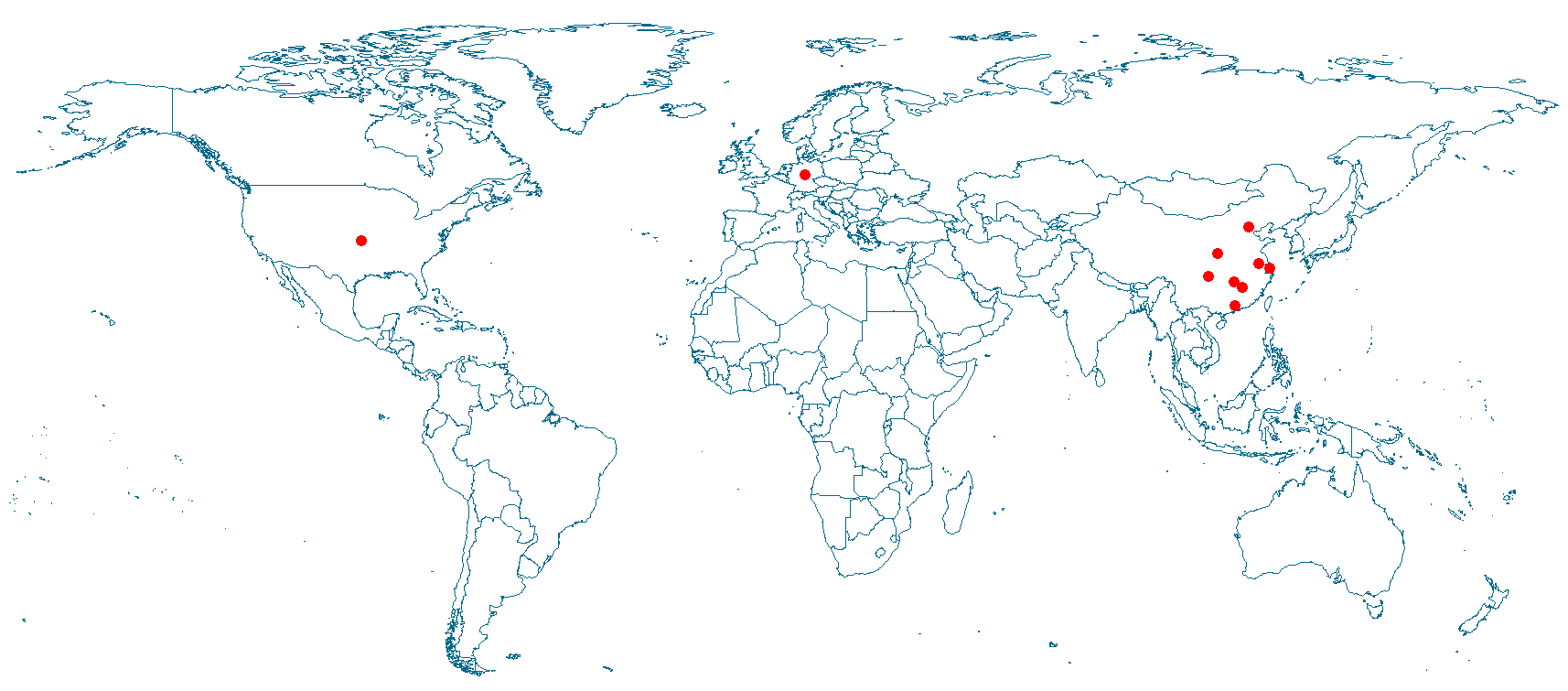
In order to better describe the nature of the images, each expert estimated the proportion of cancerous tissue in all the tissue areas. The average of their estimates was used to describe the area of the cancer tissue.

**Figure S1. Location of independent data sources indicated by red circles**

**1.b Dataset-A, B, C, and D**

The sets of images chosen from XH for Dataset-A, C, and D were mutually exclusive. We denoted them as XH-Dataset-A, XH-Dataset-C, and XH-Dataset-D accordingly. The images from other hospitals TXH, PCH, HPH, FUS, GPH, NJD, SWH, AMU, SYU, and CGH, as well as ACL were used for both patient-level testing (Dataset-C) and the Human-AI contest (Dataset-D). The Dataset-B images for patch-level validation was all from NCT-UMM. The TCGA images made from FFPE and frozen samples were used for patient-level testing only as their diagnosis is known online. The slides from different sources were distributed approximately equally in Dataset-D (Figure S2).

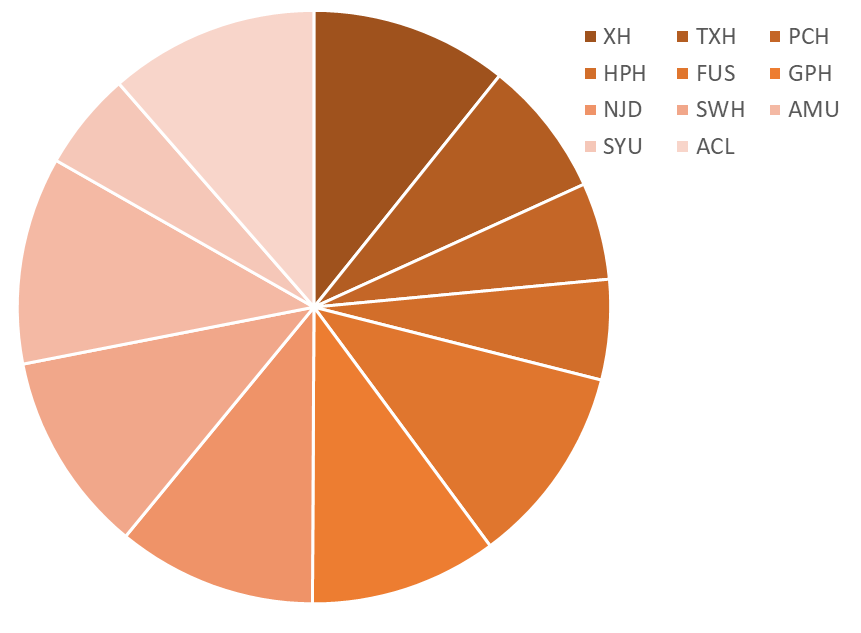
**Table S1. Allocation (number) of slides in various datasets**

| **Dataset** | **A** | **B** | **C** | **D** |
| --- | --- | --- | --- | --- |
| XH | 842 | 0 | 10003 | 213 |
| NCT-UMM | 0 | 111 | 0 | 0 |
| TXH | 0 | 0 | 135 | 135 |
| PCH | 0 | 0 | 96 | 96 |
| HPH | 0 | 0 | 99 | 99 |
| FUS | 0 | 0 | 198 | 198 |
| GPH | 0 | 0 | 185 | 185 |
| NJD | 0 | 0 | 197 | 197 |
| SWH | 0 | 0 | 199 | 199 |
| AMU | 0 | 0 | 205 | 205 |
| SYU | 0 | 0 | 97 | 97 |
| ACL | 0 | 0 | 207 | 207 |
| CGH | 0 | 0 | 100 | 0 |
| TCGA | 0 | 0 | 1793 | 0 |
| **Total** | 842 | 111 | 13,514 | 1,831 |

**Figure S2. The allocation of slides in Dataset-D from different independent sources.**

The patch-level prediction is essential for the entire approach. Since cancer is known as a heterogeneous disease, a high degree of phenotypic diversity exists “within tumors” and “among tumors”^1^. We carefully selected WSIs as Dataset-A for model training to make sure that all common tumor histological subtypes were included so that the trained model would be widely applicable to general practical diagnosis in routine clinical settings. The patches from the same patient were all put into the same data set (either training or testing) so that the training and testing data sets are independent. In total, 30,056 labeled tumor patches from 614 patients and 32,863 normal patches from 228 healthy subjects were used for model training and testing (Table 2). To ensure an appropriate and comprehensive representation of cancer and normal tissue characteristics, we included an average of 49 patches per tumor sample and 144 patches per healthy sample. We controlled the distribution of the proportion of cancer cells in patches. The number of patches containing a large proportion of cancer cells and the number of patches containing only a few cancer cells were approximately balanced so that the patches used for training were representative of cases seen in practice.

We further validated our patch-level performance using Dataset-B, which contained 107,180 patches downloaded from NCT biobank and the UMM pathology archive (NCT-UMM). There were two independent subsets: 100,000 image patches of 86 hematoxylin and eosin stain (HE) slides of human cancer tissue (NCT-CRC-HE-100K) and 7,180 image patches of 25 slides of CRC tissue (CRC-VAL-HE-7K)^2^. All images are 224x224 pixels at 0.5 microns per pixel. More description can be found at <https://zenodo.org/record/1214456#.XV2cJeg3lhF>. The patches were rescaled to default input size before they are fed to the networks for testing.

In Dataset-C, the area occupied by cancer cells varied in images from different centers. Most (~72%) of the slides from the ten hospitals and ACL contained 10%-50% cancer cells by area (Figure S3). In the contest images of Dataset-D, there are an average of ~5,045 patches on each slide, and more than 20% of the slides contain <1000 patches.

After the slides were digitized, the visual verification of the cancer diagnosis labels was performed with high stringency and accuracy. Dataset-A and Dataset-C included more than 10,000 slides, which were independently reviewed by two senior and seasoned pathologists with initial and second read. When their diagnoses were consistent with the previous clinical diagnosis conclusion, the slides were then included in the dataset. If the two experts disagreed with each other or with the previous clinical diagnosis, the slides were excluded. The labels of slides from TCGA were obtained from the original TCGA database. The labels of Dataset-B were from the NCT-UMM. The binary labels of Dataset-D for the Human-AI contest were more strictly checked. Three senior highly experienced pathologists independently reviewed the pathological images without knowing the previous clinical diagnosis. If a consensus was reached, the slides were included; otherwise, two other independent pathologists would join the review. After a discussion among the five pathologists, the sample was included only if they reached an agreement; otherwise, it was excluded.

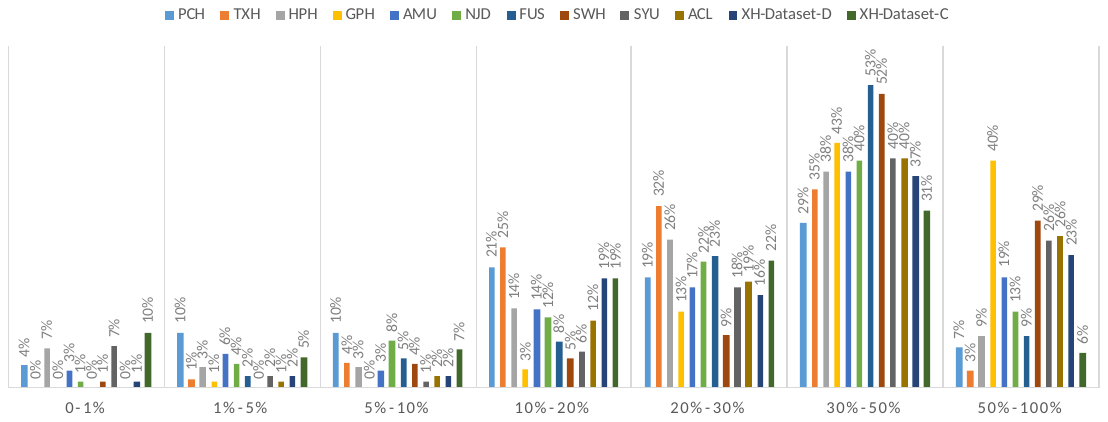

| P | PCH | TXH | HPH | GPH | AMU | NJD | FUS | SWH | SYU | ACL | XH-Dataset-D | XH-Dataset-C |
| --- | --- | --- | --- | --- | --- | --- | --- | --- | --- | --- | --- | --- |
| 0-1% | 4% | 0% | 7% | 0% | 3% | 1% | 0% | 1% | 7% | 0% | 1% | 10% |
| 1%-5% | 10% | 1% | 3% | 1% | 6% | 4% | 2% | 0% | 2% | 1% | 2% | 5% |
| 5%-10% | 10% | 4% | 3% | 0% | 3% | 8% | 5% | 4% | 1% | 2% | 2% | 7% |
| 10%-20% | 21% | 25% | 14% | 3% | 14% | 12% | 8% | 5% | 6% | 12% | 19% | 19% |
| 20%-30% | 19% | 32% | 26% | 13% | 17% | 22% | 23% | 9% | 18% | 19% | 16% | 22% |
| 30%-50% | 29% | 35% | 38% | 43% | 38% | 40% | 53% | 52% | 40% | 40% | 37% | 31% |
| 50%-100% | 7% | 3% | 9% | 40% | 19% | 13% | 9% | 29% | 26% | 26% | 23% | 6% |

**Figure S3. The distribution of cancerous area in multiple independent WSI datasets measured by the proportion of patches (P) containing cancer cells on the WSI. The estimate of proportions was provided by the pathologist.**

**1.c Comparison of different CNN architectures at patch- and patient-level**

In choosing the architecture for our Convolutional Neural Network (CNN), we tested and compared the relative performances in our datasets several architectures that are commonly used in histological and other image analyses, including: VGGNet^3^, Inception-v1^4^, Inception-v3^5^, and Inception-v4 (ResNet-v2)^6^. We compared their relative performances at patch-level (XH-Dataset-A, NCT-CRC-HE-100K in Dataset-B), patient-level (XH-Dataset-C, PCH, TXH, HPH, FUS, GPH, SWH, SYU-CGH, AMU, NJD, ACL, TCGA), and using Human-AI contest testing datasets (XH-Dataset-D, PCH, TXH, HPH, ACL). A cluster size of 4 was used for the patient-level comparison.

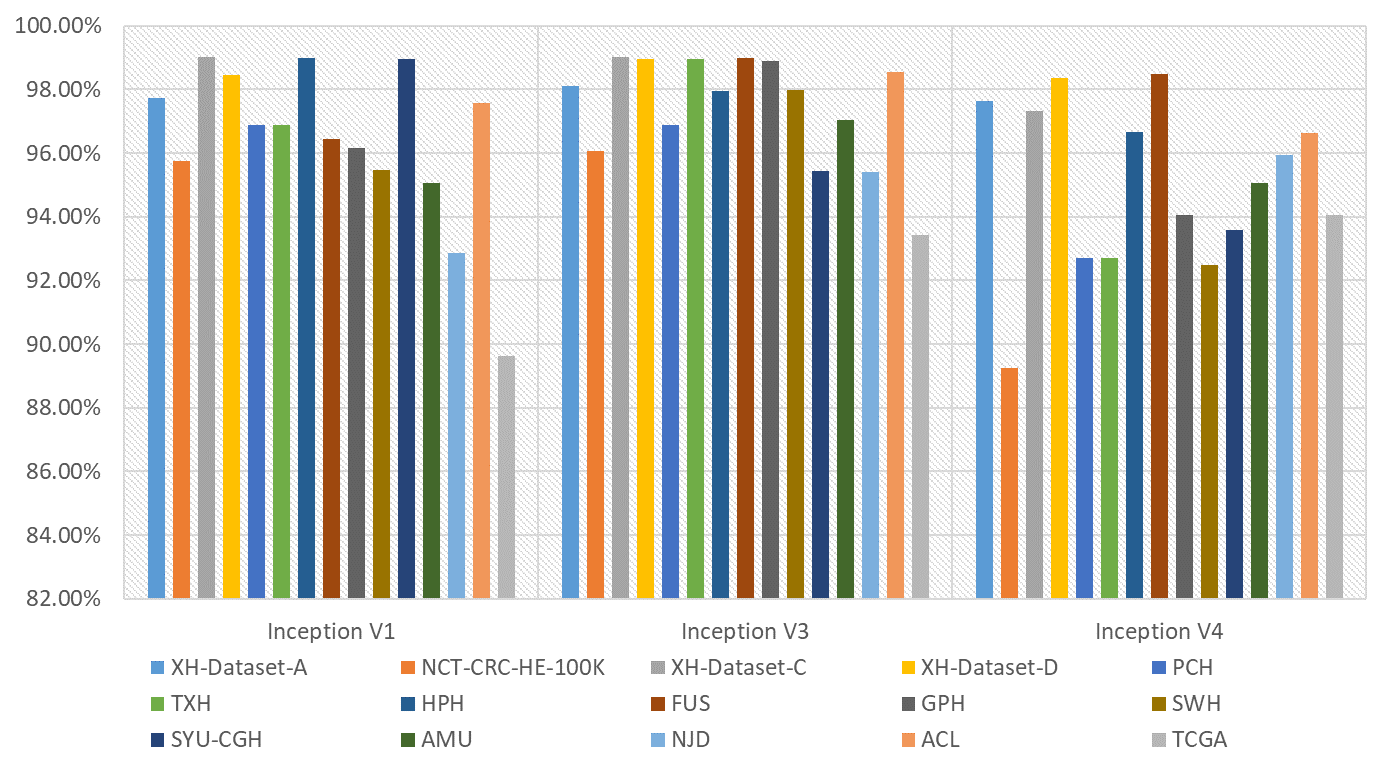
Inception-v3 was finally selected to be the architecture used in our approach. In Table S2, Inception-v3 yielded the highest accuracy and AUC using patch-level XH-Dataset-A and patient-level XH-Dataset-B, which was the largest dataset by sample size. In Figure S4, the accuracies of the Inception-v3 network were high and stable for almost all the datasets relative to Inception-v1 and Inception-v4. The only performance deficiency of Inception-v3 was on the TCGA-Frozen dataset, which may be explained by the different preparation with frozen samples and/or other sample preprocessing procedures that are different from the data sets used for training. The result from VGGNet was worse than the three Inception CNN architectures and thus not shown here. Therefore, we choose Inception-V3 as the architecture of the CNN model in our approach.

**Figure S4. Accuracies of Inception architectures on the datasets**

**Table S2. Relative performance of Google Inception architectures on the datasets**

| Architecture | Data source | Sensitivity | Specificity | Accuracy | AUC |
| --- | --- | --- | --- | --- | --- |
| Inception V1 | XH-Dataset-A | 95.33% | 99.31% | 97.72% | 99.77% |
|  | NCT-CRC-HE-100K | 94.31% | 95.98% | 95.74% | 98.66% |
|  | XH-Dataset-C | 98.80% | 99.51% | 99.02% | 99.16% |
|  | XH-Dataset-D | 97.96% | 98.97% | 98.46% | 98.46% |
|  | PCH | 100% | 93.48% | 96.88% | 96.74% |
|  | TXH | 98.00% | 95.74% | 96.88% | 96.85% |
|  | HPH | 100% | 97.96% | 98.98% | 98.98% |
|  | FUS | 100% | 92.86% | 96.46% | 97.96% |
|  | GPH | 100% | 91.76% | 96.17% | 96.47% |
|  | SWH | 98.99% | 92.00% | 95.48% | 98.43% |
|  | SYU-CGH | 100.00% | 83.33% | 98.97% | 91.67% |
|  | AMU | 100% | 90.20% | 95.07% | 97.55% |
|  | NJD | 95.96% | 89.69% | 92.86% | 95.71% |
|  | ACL | 99% | 96.26% | 97.58% | 97.63% |
|  | TCGA-Frozen | 89.49% | 91.04% | 89.64% | 95.45% |
| Inception V3 | XH-Dataset-A | 96.99% | 99.22% | 98.11% | 99.83% |
|  | NCT-CRC-HE-100K | 92.03% | 96.74% | 96.07% | 98.32% |
|  | XH-Dataset-C | 98.80% | 99.51% | 99.02% | 99.16% |
|  | XH-Dataset-D | 97.96% | 100.00% | 98.97% | 98.98% |
|  | PCH | 96.00% | 97.83% | 96.88% | 96.91% |
|  | TXH | 100.00% | 97.92% | 98.96% | 98.96% |
|  | HPH | 97.96% | 97.96% | 97.96% | 97.96% |
|  | FUS | 100% | 97.96% | 98.99% | 98.98% |
|  | GPH | 100% | 97.65% | 98.91% | 98.82% |
|  | SWH | 98.99% | 97.00% | 97.99% | 97.99% |
|  | SYU-CGH | 98.90% | 92.45% | 95.43% | 95.68% |
|  | AMU | 97% | 97.06% | 97.04% | 97.04% |
|  | NJD | 92.93% | 97.94% | 95.41% | 95.43% |
|  | ACL | 100% | 97.20% | 98.55% | 98.60% |
|  | TCGA-Frozen | 94.04% | 88.06% | 93.44% | 91.05% |
| Inception V4 | XH-Dataset-A | 96.95% | 98.34% | 97.64% | 97.81% |
|  | NCT-CRC-HE-100K | 94.08% | 88.42% | 89.24% | 97.10% |
|  | XH-Dataset-C | 99.33% | 96.97% | 97.32% | 98.14% |
|  | XH-Dataset-D | 97.67% | 98.97% | 98.36% | 98.32% |
|  | PCH | 98.00% | 86.96% | 92.71% | 92.48% |
|  | TXH | 100.00% | 85.42% | 92.71% | 92.71% |
|  | HPH | 100.00% | 93.33% | 96.67% | 96.67% |
|  | FUS | 100% | 96.94% | 98.48% | 98.50% |
|  | GPH | 100% | 87.06% | 94.05% | 95.70% |
|  | SWH | 100% | 85.00% | 92.50% | 93.20% |
|  | SYU-CGH | 100% | 87.46% | 93.58% | 94.10% |
|  | AMU | 100% | 90.20% | 95.07% | 96.50% |
|  | NJD | 99.00% | 92.78% | 95.93% | 95.90% |
|  | ACL | 96.00% | 97.20% | 96.62% | 96.60% |
|  | TCGA-Frozen | 93.66% | 96.36% | 94.06% | 95.01% |

**1.d Image preprocessing and CNN training at patch-level**

There were 3 steps in the image preprocessing process. First, we tiled each WSI at 20× magnification with non-overlapping 300×300 pixel patches, which can be easily transformed to the required input size of most CNN architectures (such as the 299×299 input size required by Inception-v3^5^, Table S3). The use of a smaller patch size compared with other studies with patches of 512×512 pixels would make the boundaries of cancer regions more accurate^5^^;^ ^25^. Second, we removed non-informative background patches according to two criteria: the maximum difference among the 3 color channel values of the patch was less than 20, or the brightness of more than 50% of the patch surface was less than 220 in grayscale^7^. Combining these two criteria, we removed background patches and kept as many tissue patches as possible. Third, regular image augmentation procedures were applied, such as random flipping and random adjustment of the saturation, brightness, contrast, and hue. The color of each pixel was centered by the mean of each image and its range was converted/normalized from [0, 255] to [-1, 1].

**Table S3. Input patch size for common CNN**

|  | Patch size | Notes |
| --- | --- | --- |
| Patches in training sets | 300*300*3 | The size of Labeled patch by Pathologists |
| Inception V1 | 224*224*3 | Default input size |
| Inception V3 | 299*299*3 | Default input size |
| Inception V4 | 299*299*3 | Default input size |
| VGG 19 | 224*224*3 | Default input size |
| ResNet-101 | 224*224*3 | Default input size |

We initialized the network by loading pre-trained weights achieved in ImageNet^5^. The 300×300 pixel patches were resized to a size of 299×299 pixels. Accordingly, the patches in the testing sets were rescaled to 299×299 pixels (0.37 μm/pixels) before they were fed to the network. The network was deeply fine-tuned by following training steps. Given the possible high false positive rate after aggregating the patch-level results, the optimal set of hyper-parameters was randomly searched with an objective of reaching >95% sensitivity and >99% specificity. We showed that, with this objective at the patch-level, the error rate at the patient-level was well controlled. We define the following metrics used in our evaluations: sensitivity (Se) = (True predicted positives)/(Real positives), specificity (Sp) = (True predicted negatives)/(Real negatives), and false positive rate (FPR) = (False predicted positives)/(Real negatives) = 1- Sp. The sensitivity measures how the model can diagnose real colorectal cancer (CRC) patients as CRC cases, while the specificity measures how the model can identify non-CRC patients as non-CRC subjects. Low sensitivity will lead to misdiagnosis and no treatment of CRC patients. Low specificity will lead to misdiagnosis and wrong treatment of non-CRC subjects.

Our approach for patient diagnosis was based on the aggregation of patch-level predictions. The performance of patch-level prediction would determine the accuracy of patient-level diagnosis. Assuming the patch-level sensitivity and specificity were $\theta$ and $\gamma$, the cluster size for aggregation was k, and each patient had only one histological image slide, if the input patches contained complete cancer or non-cancer information and were mutually independent, the theoretical probability of correctly identifying CRC patients (patient-level Se) was $\theta^{k}$, and the probability of falsely identifying non-CRC patients (patient-level FPR) was ${(1-\gamma)}^{k}$. When $k=3$, patch-level Sp=0.95, we have a patch-level FPR=0.05, while the patient-level FPR≈0.0001. The above theoretical expectation is based on the assumption that each patch on the same WSI is independent. In practice, the nearby patches were highly correlated; consequently, the theoretical derivation was not valid for our aggregation result. However, the empirical results, including examples in Table S2, showed that a patch-level sensitivity of ~95% and specificity of ~99% was sufficient to achieve a high predictive power and control the FPR at the patient-level.

The network was finalized after 150,000 epochs of fine-tuning the parameters at all layers using the RMSProp^8^ optimizer with a weight decay of 0.00004, a momentum value of 0.9, and RMSProp decay set to 0.9. The initial learning rate was 0.01 and was exponentially decayed with epochs to the final learning rate of 0.0001. The optimized result was achieved when the batch size was 64. The training and testing procedures were implemented in a Linux server with an NVIDIA P100 GPU. We used Python v2.7.15 and Tensorflow v1.8.0 for data preprocessing and CNN model training and testing.

**1.e Comparison of different cluster sizes for aggregation of patch-level results for patient level diagnosis**

We compared the performance of different cluster sizes on aggregating the patch-level results to patient-level prediction. The patient-level testing datasets were used for the comparison, including XH-Dataset-C, XH-Dataset-D, PCH, TXH, HPH, FUS, GPH, NJD, SWH, AMU, SYU, CGH, ACL, and frozen TCGA samples.

We selected the cluster size of 4 in our approach because it resulted in the highest accuracy in most of the datasets (except the TCGA), which also represented the best balance between sensitivity and specificity. Using the results from XH-Dataset-C as an illustrating example, the cluster size of four was able to achieve high sensitivity (98.80%) and specificity (99.51%), while the size of three yielded slightly higher sensitivity (99.52%) and lower specificity (97.78%). The cluster size of 2 was good with respect to the diagnosis rate of CRC (sensitivity 99.72%). Generally, increasing the cluster size would lead to higher specificity but lower sensitivity, which was illustrated in Figure S5 using the largest test set of XH-Dataset-C.

**Table S4. Comparison of cluster size for aggregation of patch-level results**

| Cluster Size | Data source | Sensitivity | Specificity | Accuracy | AUC |
| --- | --- | --- | --- | --- | --- |
| 2 continuous patches | XH-Dataset-C | 99.72% | 93.67% | 97.81% | 96.70% |
|  | XH-Dataset-D | 100% | 87.63% | 93.85% | 93.81% |
|  | PCH | 100% | 71.74% | 86.46% | 85.87% |
|  | TXH | 100% | 72.92% | 86.46% | 86.46% |
|  | HPH | 100% | 73.47% | 86.73% | 86.73% |
|  | FUS | 100% | 66.33% | 83.33% | 83.16% |
|  | GPH | 100% | 82.35% | 91.80% | 91.18% |
|  | NJD | 98.99% | 62.89% | 81.12% | 80.94% |
|  | SWH | 100% | 61.00% | 80.40% | 80.50% |
|  | AMU | 100% | 77.45% | 88.67% | 88.73% |
|  | SYU-CGH | 100% | 60.38% | 78.68% | 80.19% |
|  | ACL | 100% | 74.77% | 86.96% | 87.38% |
|  | TCGA-Frozen | 98.43% | 55.22% | 94.11% | 76.83% |
| 3 continuous patches | XH-Dataset-C | 99.52% | 97.78% | 98.97% | 98.65% |
|  | XH-Dataset-D | 98.98% | 93.81% | 96.41% | 96.40% |
|  | PCH | 100% | 91.30% | 95.83% | 95.65% |
|  | TXH | 100% | 83.33% | 91.67% | 91.67% |
|  | HPH | 100% | 87.76% | 93.88% | 93.88% |
|  | FUS | 100% | 90% | 94.95% | 94.90% |
|  | GPH | 100% | 92.94% | 96.72% | 96.47% |
|  | NJD | 95.96% | 85.57% | 90.82% | 90.76% |
|  | SWH | 100% | 84.00% | 91.96% | 92.00% |
|  | AMU | 99% | 89.22% | 94.09% | 94.11% |
|  | SYU-CGH | 100% | 81.13% | 89.85% | 90.57% |
|  | ACL | 100% | 89.72% | 94.69% | 94.86% |
|  | TCGA-Frozen | 97.19% | 82.84% | 95.75% | 90.01% |
| 4 continuous patches | XH-Dataset-C | 98.80% | 99.51% | 99.02% | 99.16% |
|  | XH-Dataset-D | 97.96% | 100.00% | 98.97% | 98.98% |
|  | PCH | 96.00% | 97.83% | 96.88% | 96.91% |
|  | TXH | 100.00% | 97.92% | 98.96% | 98.96% |
|  | HPH | 97.96% | 97.96% | 97.96% | 97.96% |
|  | FUS | 100% | 97.96% | 98.99% | 98.98% |
|  | GPH | 100% | 97.65% | 98.91% | 98.82% |
|  | NJD | 92.93% | 97.94% | 95.41% | 95.43% |
|  | SWH | 98.99% | 97.00% | 97.99% | 97.99% |
|  | AMU | 97% | 97.06% | 97.04% | 97.04% |
|  | SYU-CGH | 98.90% | 92.45% | 95.43% | 95.68% |
|  | ACL | 100% | 97.20% | 98.55% | 98.60% |
|  | TCGA-Frozen | 94.04% | 88.06% | 93.44% | 91.05% |

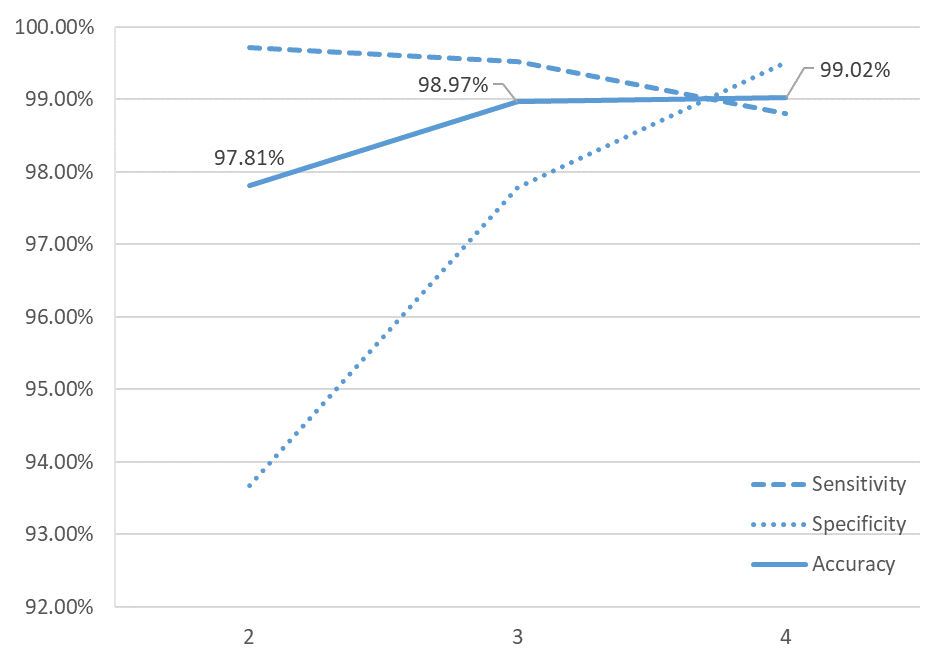
**Figure S5. Performance change with the increase of the cluster size (x-axis) on XH-Dataset-C**

**1.f Statistical analysis and visualization**

For a patient who had one or multiple slides, denoted by $S=\{s_{1}, s_{2},\ldots,s_{l}\}$, we provided the patient-level diagnosis $D\left( S \right)$ combining the results from all of his/her slides: $D\left( S \right)=D\left( s_{1} \right)\cup D\left( s_{2} \right)\cup\ldots\cup D(s_{l})$, where $D\left( s_{l} \right)=$1 or 0 indicated a positive or negative classification of the $l$-th slide respectively. So that the patient will be diagnosed as having cancer as long as one of the slides is diagnosed so.

We assessed the performance of the AI and pathologists in terms of sensitivity, specificity, and accuracy ($\frac{\# of correct predictions}{\# of total predictions}$) for the diagnosis. The receiver operating characteristic (ROC) curve that plotted the sensitivity versus the FPR and the corresponding area under the ROC curve (AUC) was computed. We examined the pairwise agreements among AI and pathologists by the Cohen’s Kappa statistic ($K$). The statistical analyses were done in R (Vienna, Austria). To locate the CRC region in the WSI, we visualized the WSI as a heatmap based on the confidence score of each patch. The brighter a region, the more confident the classifier would consider that region cancer positive.

**Supplementary-Table 1. Pathologist info**

| **Pathologist ID** | **Years in Clinic** | **Title** |
| --- | --- | --- |
| A | 1 | Resident physician |
| B | 3 | Resident physician |
| C | 5 | Physician-in-charge |
| D | 7 | Physician-in-charge |
| E | 12 | Physician-in-charge |
| F | 18 | Associate chief physician |

**Supplementary-Table** 2**. List of AUCs of AI applied in CRC and other cancer types**

| Study | Patch-level test data | | Independent patch-level test data | | Slide-level test data | | Independent slide-level test data | | |
| --- | --- | --- | --- | --- | --- | --- | --- | --- | --- |
|  | Number (#) of patches | AUC | # of patches | AUC | # of slides | AUC | # of datasets | # of slides | AUC |
| **Colorectal cancer** | | | | | | | | | |
| Haj‑Hassan et al.^9^ | NA | Unsegmented~0.7923  Segmented~0.9917 | NA | NA | NA | NA | NA | NA | NA |
| Xu et al.^10^ | 717 | 0.969-0.980^a^ | NA | NA | NA | NA | NA | NA | NA |
| Sari et al.^11^ | 1,592 | 0.994 | NA | NA | NA | NA | NA | NA | NA |
| Kainz et al.^12^ | 60 | 0.983^a^ | 20^a^ | 0.950^a^ | NA | NA | NA | NA | NA |
| Kather et al.^2^ | 100,000 | 0.987 | 7,180 | 0.943 | NA | NA | NA | NA | NA |
| Ponzio et al.^13^ | 4500 | 0.9037-0.9682 | NA | NA | NA | NA | NA | NA | NA |
| **Other cancers** | | | | | | | | | |
| Coudray et al.^14^ | NA | NA | NA | NA | 244 | 0.990-0.993 | 3 | 340 | LUAD~0.833-0.913  LUSC~0.861-0.941 |
| Cruz-Roa et al.^15^ | 50,963 | 0.842^b^ | NA | NA | NA | NA | NA | NA | NA |
| Araujo et al.^16^ | 240 | 0.829^a^ | 192 | 0.693^a^ | 20 | 0.900^a^ | 1 | 16 | 0.750^a^ |
| Motlagh et al.^17^ | 2,147 | 0.999 | NA | NA | NA | NA | NA | NA | NA |
| Campanella et al.^18^ | NA | NA | NA | NA | 12,132 | 0.986-0.991 | 1 | 12,727 | 0.986-0.991 |
| Campanella et al.^18^ | NA | NA | NA | NA | 6,252 | 0.986-0.988 | 1 | 3,710 | 0.986-0.988 |
| Campanella et al.^18^ | NA | NA | NA | NA | 8,670 | 0.965-0.966 | 1 | 1,224 | 0.965-0.966 |
| **Our study** | 20,264 | 0.998 | 107,180 | 0.983-0.985 | 10,003 | 0.992 | 12 | 3,065 | 0.911-0.992^c^ |

Note: a: accuracy; b: balanced accuracy; c, aggregated on 4 continuous patches

**Supplementary-Table** 3**. Patient-level results** **on Dataset-C & D**

| **Source** | **Sensitivity** | **Specificity** | **Accuracy** | **AUC** |
| --- | --- | --- | --- | --- |
| **Dataset-C only** | | | | |
| XH | 98.80% | 99.51% | 99.02% | 99.16% |
| TCGA-Frozen | 94.04% | 88.06% | 93.44% | 91.05% |
| TCGA-FFPE | 97.96% | 100.00% | 97.98% | 98.98% |
| SYU-CGH | 98.90% | 92.45% | 95.43% | 95.68% |
| **Dataset-D (Human AI contest) only** | | | | |
| XH | 97.96% | 100% | 98.97% | 98.98% |
| SYU | 98.90% | 100% | 98.97% | 99.45% |
| **Dataset-C & D** | | | | |
| PCH | 96.00% | 97.83% | 96.88% | 96.91% |
| TXH | 100% | 97.92% | 98.96% | 98.96% |
| HPH | 97.96% | 97.96% | 97.96% | 97.96% |
| FUS | 100% | 97.96% | 98.99% | 98.98% |
| GPH | 100% | 97.65% | 98.91% | 98.82% |
| NJD | 92.93% | 97.94% | 95.41% | 95.43% |
| SWH | 98.99% | 97.00% | 97.99% | 97.99% |
| AMU | 97% | 97.06% | 97.04% | 97.04% |
| ACL | 100% | 97.20% | 98.55% | 98.60% |

**Supplementary-Table 4. Overall performance of AI and pathologists in Human-AI contest**

|  | AI | Pathologists | | | | | | |
| --- | --- | --- | --- | --- | --- | --- | --- | --- |
|  |  | A | B | C | D | E | F | Average |
| Sensitivity | 98.16% | 98.08% | 97.26% | 96.71% | 98.26% | 98.53% | 96.00% | 97.47% |
| Specificity | 98.05% | 92.19% | 94.87% | 98.90% | 99.09% | 98.17% | 95.26% | 96.41% |
| Accuracy | 98.06% | 95.81% | 96.73% | 97.70% | 98.56% | 98.38% | 95.64% | 97.14% |
| AUC | 98.10% | 95.14% | 96.07% | 97.83% | 98.67% | 98.35% | 95.63% | 96.95% |

**Supplementary-Table 5. Cohen's kappa coefficient for agreement among Human experts and AI**

|  | Human Experts | | | | | |
| --- | --- | --- | --- | --- | --- | --- |
|  | A | B | C | D | E | F |
| AI | 0.891 | 0.896 | 0.905 | 0.919 | 0.908 | 0.858 |
| A |  | 0.931 | 0.924 | 0.920 | 0.920 | 0.813 |
| B |  |  | 0.938 | 0.935 | 0.921 | 0.841 |
| C |  |  |  | 0.944 | 0.928 | 0.851 |
| D |  |  |  |  | 0.945 | 0.880 |
| E |  |  |  |  |  | 0.851 |

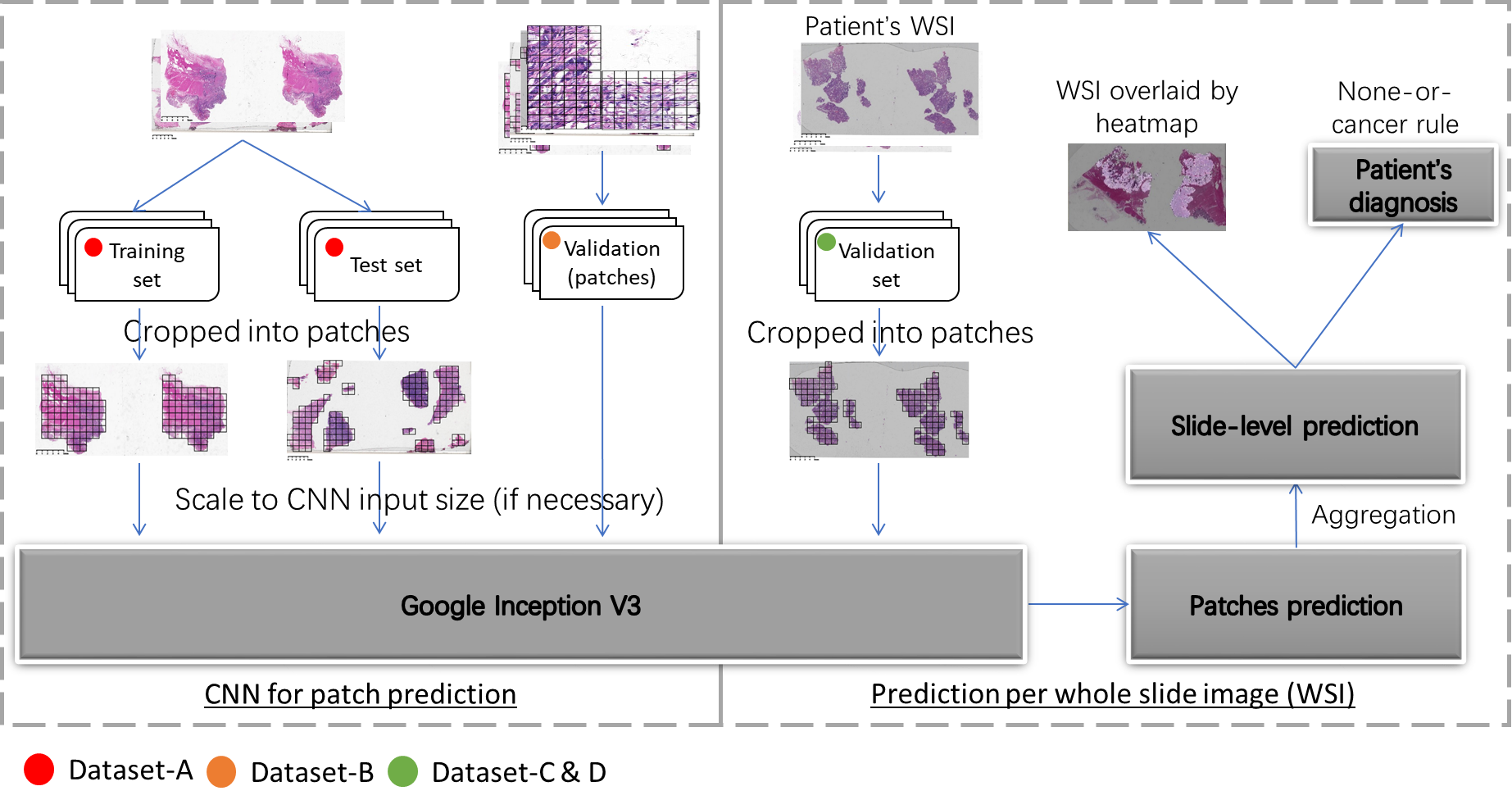
**Supplementary-Figure 1. Study pipeline**

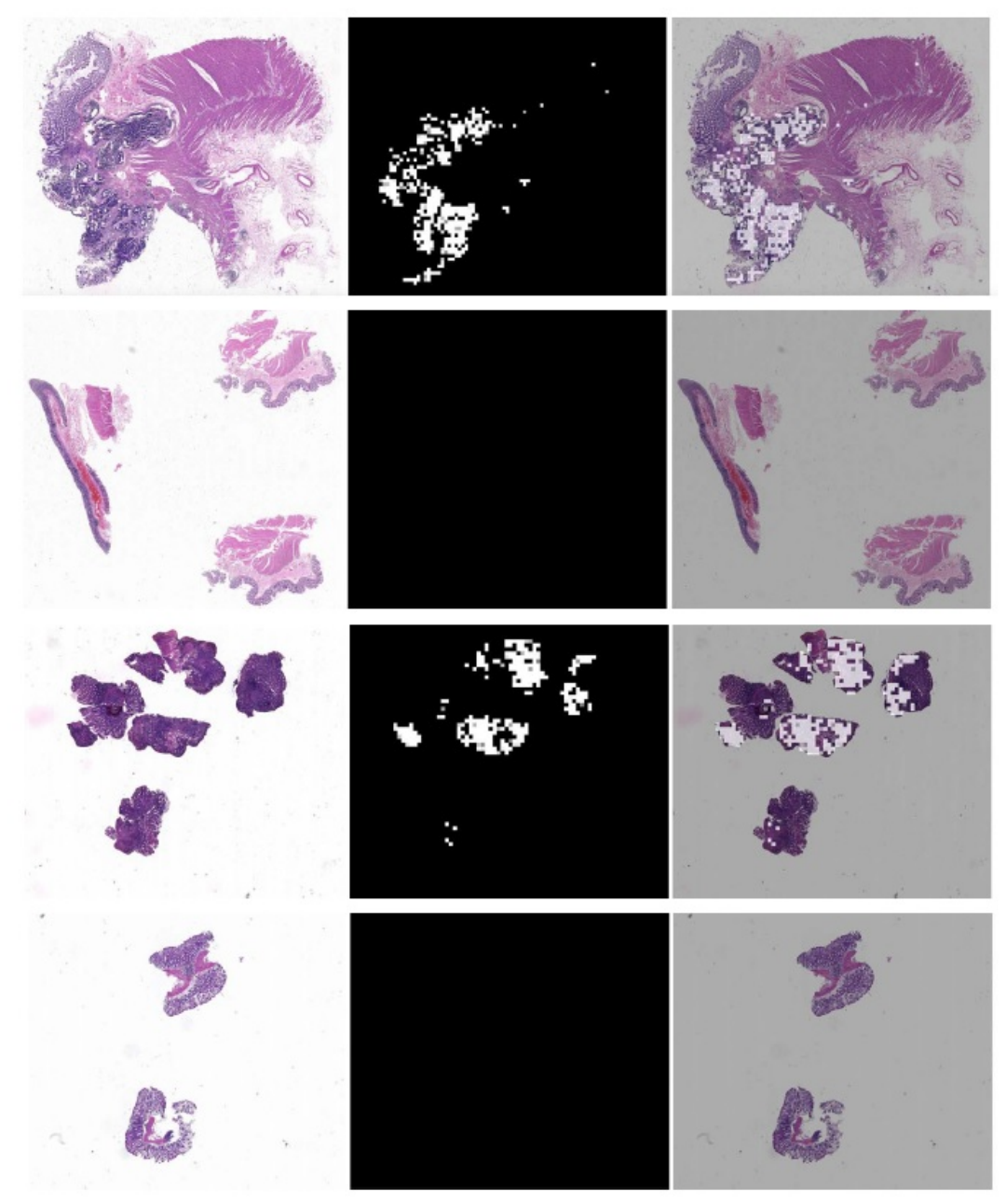
**Supplementary-Figure 2. Heatmap produced by AI. Left column – WSI, middle column – predicted heatmap, right column – heatmap overlaid on the WSI. Top row: positive case from radical surgery; second row: negative case from radical surgery; third row: positive case colonoscopy; bottom row: negative case colonoscopy.**

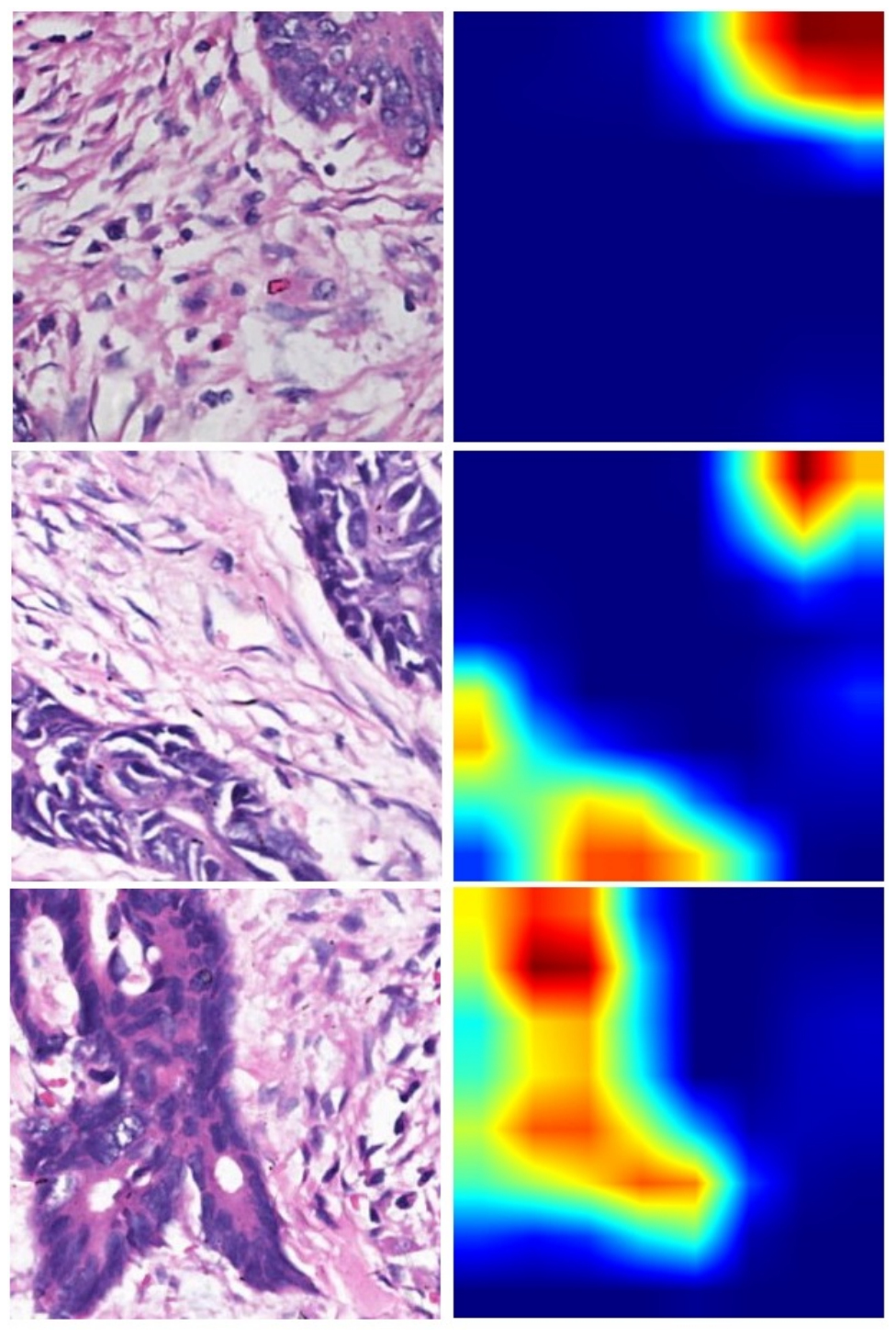
**Supplementary-Figure 3. Activation map produced by AI. Left column – WSI, right column – activation map. Each row is a pair of WSI and activation map from the same patch. The heat color indicates informative regions used by the DL AI for CRC detection.**

**Reference**

1.	Mimori K, Saito T, Niida A, Miyano S. Cancer evolution and heterogeneity. Annals of gastroenterological surgery 2018;2:332-8.

2.	Kather JN, Krisam J. Predicting survival from colorectal cancer histology slides using deep learning: A retrospective multicenter study. 2019;16:e1002730.

3.	Simonyan K, Zisserman A. Very deep convolutional networks for large-scale image recognition. arXiv preprint arXiv:14091556 2014.

4.	Szegedy C, Wei L, Yangqing J, et al. Going deeper with convolutions. 2015 IEEE Conference on Computer Vision and Pattern Recognition (CVPR); 2015 7-12 June 2015. p. 1-9.

5.	Szegedy C, Vanhoucke V, Ioffe S, Shlens J, Wojna Z. Rethinking the Inception Architecture for Computer Vision. 2016 IEEE Conference on Computer Vision and Pattern Recognition (CVPR); 2016 27-30 June 2016. p. 2818-26.

6.	Szegedy C, Ioffe S, Vanhoucke V, Alemi AA. Inception-v4, inception-resnet and the impact of residual connections on learning. Thirty-First AAAI Conference on Artificial Intelligence; 2017.

7.	Coudray N, Ocampo PS, Sakellaropoulos T, et al. Classification and mutation prediction from non-small cell lung cancer histopathology images using deep learning. Nat Med 2018.

8.	Hinton GS, N; Swersky, K. Lecture 6a Overview of Mini-Batch Gradient Descent. Lecture Notes Distributed in CSC321 of University of Toronto. 2014.

9.	Haj-Hassan H, Chaddad A, Harkouss Y, Desrosiers C, Toews M, Tanougast C. Classifications of Multispectral Colorectal Cancer Tissues Using Convolution Neural Network. J Pathol Inform 2017;8:1.

10.	Xu Y, Jia Z, Wang LB, et al. Large scale tissue histopathology image classification, segmentation, and visualization via deep convolutional activation features. BMC Bioinformatics 2017;18:281.

11.	Sari CT, Gunduz-Demir C. Unsupervised Feature Extraction via Deep Learning for Histopathological Classification of Colon Tissue Images. IEEE transactions on medical imaging 2018.

12.	Kainz P, Pfeiffer M, Urschler M. Segmentation and classification of colon glands with deep convolutional neural networks and total variation regularization. PeerJ 2017;5:e3874.

13.	Ponzio F, Macii E, Ficarra E, Di Cataldo S. Colorectal Cancer Classification using Deep Convolutional Networks - An Experimental Study 2018.

14.	Coudray N, Ocampo PS, Sakellaropoulos T, et al. Classification and mutation prediction from non-small cell lung cancer histopathology images using deep learning. Nature Medicine 2018;24:1559-67.

15.	Gurcan MN, Madabhushi A, Cruz-Roa A, et al. Automatic detection of invasive ductal carcinoma in whole slide images with convolutional neural networks. 2014;9041:904103.

16.	Araujo T, Aresta G, Castro E, et al. Classification of breast cancer histology images using Convolutional Neural Networks. PloS one 2017;12:e0177544.

17.	Jannesari M, Habibzadeh M, Aboulkheyr H, et al. Breast Cancer Histopathological Image Classification: A Deep Learning Approach. 2018 IEEE International Conference on Bioinformatics and Biomedicine (BIBM); 2018 3-6 Dec. 2018. p. 2405-12.

18.	Campanella G, Hanna MG, Geneslaw L, et al. Clinical-grade computational pathology using weakly supervised deep learning on whole slide images. Nature Medicine 2019.
